# Rapid Whole Cell Imaging Reveals A Calcium-APPL1-Dynein Nexus That Regulates Cohort Trafficking of Stimulated EGF Receptors

**DOI:** 10.1101/481796

**Authors:** H M York, A Patil, U K Moorthi, A Kaur, A Bhowmik, G J Hyde, H Gandhi, K Gaus, S Arumugam

## Abstract

Multicellular life processes such as proliferation and differentiation depend on cell surface signaling receptors that bind ligands generally referred to as growth factors. Recently, it has emerged that the endosomal system provides rich signal processing capabilities for responses elicited by these factors [1-3]. At the single cell level, endosomal trafficking becomes a critical component of signal processing, as exemplified by the epidermal growth factor (EGF) receptors of the receptor tyrosine kinase family. EGFRs, once activated by EGF, are robustly trafficked to the phosphatase-enriched peri-nuclear region (PNR), where they are dephosphorylated [4-8]. However, the details of the mechanisms regulating the movements of stimulated EGFR in time and space, i.e., towards the PNR, are not known. What endosomal regulators provide specificity to EGFR? Do modifications to the receptor upon stimulation regulate its trafficking? To understand the events leading to EGFR translocation, and especially the early endosomal dynamics that immediately follow EGFR internalization, requires the real-time, long-term, whole-cell imaging of multiple elements. Here, exploiting the advantages of lattice light-sheet microscopy [9], we show that the binding of EGF by its receptor, EGFR, triggers a transient calcium increase that peaks by 30 s, causing the desorption of APPL1 from pre-existing endosomes within one minute, the rebinding of liberated APPL1 to EGFR within three minutes, and the dynein-dependent translocation of APPL1-EGF-bearing endosomes to the PNR within five minutes. The novel, cell spanning, fast acting network that we reveal integrates a cascade of events dedicated to the cohort movement of activated EGFR receptors. Our findings support the intriguing proposal that certain endosomal pathways have shed some of the stochastic strategies of traditional trafficking, and have evolved behaviors whose predictability is better suited to signaling [10, 11]. Work presented here demonstrates that our whole cell imaging approach can be a powerful tool in revealing critical transient interactions in key cellular processes such as receptor trafficking.

## INTRODUCTION

The endocytic system is critical to a cell’s ability to faithfully transduce signals and to process their extracellular environment. At the plasma membrane, EGF stimulation results in multiple parallel processes — transient increases in Ca^2+^ [12, 13], rapid reorganization of actin filaments [14] and formation of new clathrin-coated pits (CCPs) [15]. EGFR has been compared to transferrin in many studies as an example of a ligand-induced, rather than a constitutive receptor, system[16, 17]. Stimulated EGFRs initiate new CCPs that are distinct from transferrin receptor containing CCPs [15, 18, 19]. Post internalization, the intracellular itineraries of the two receptors are distinct with stimulated EGFR localizing to late endosomes and transferrin receptors recycled back through the early endosomes marked by Rab5[16].

Rab5 effectors such as EEA1 and APPL1 and 2 have been shown to be involved in the pre-early endosomal steps of endosomal maturation [20]. APPL1 can bind directly to Rab5 [21] as well as to a lipid bilayer via a PH BAR domain [22]. APPL1 has been also shown to bind the cytosolic tail of various receptors, including the adiponectin receptor [23], the nerve growth factor receptor [24] and EGFR [20, 25] via its PTB domain. Following endocytosis, EGFR has been shown to enter a population of endosomes marked by APPL1 [10, 20, 26]. It has also been demonstrated that endosomes bearing a subset of clathrin dependent cargoes, including EGFR, are more mobile and mature faster post internalization [27], compared to those that carry transferrin. To better understand the apparent central role of APPL1 in regulating EGFR, its relation to the observed rapid dynamics of EGF bearing endosomes, and more generally, the molecular events and processes that lead to intracellular divergence of EGFR and transferrin receptors, we measure the dynamics of the newly formed EGF bearing endosomes as well as APPL1 within the first ten minutes of stimulation by EGF. Capitalizing on the rapid volumetric imaging capabilities of Lattice Light-Sheet Microscopy (LLSM), we visualized the first events of internalization of EGF stimulated receptors and discovered that 1. APPL1 relocates from pre-existing endosomes onto EGFR puncta and 2. APPL1 binding to EGFR results in a cohort, dynein mediated movement of EGFR towards the peri-nuclear region of the cell where the endosomal maturation and signal quenching of EGFRs occur. The presented novel behavior of EGF bearing endosomes is a significant detail in understanding signal processing by live cells that has not been described elsewhere to our knowledge.

## RESULTS

### APPL1 responds to EGF stimulation by binding immediately to EGFR bearing endosomes and display enhanced retrograde motility

To investigate the whole cell distribution of APPL1 on receptor bearing endosomes following internalization of fluorescently labeled EGF or transferrin in living HeLa cells transfected with APPL1-eGFP, we took advantage of the more rapid volumetric imaging and decreased photo-bleaching offered by a lattice light-sheet microscope [9] (LLSM). Fast multi-color, whole-cell imaging revealed the temporal dynamics of all fluorescently labeled endosomal populations. To capture the first cargo bearing endosomes post-internalization with a time resolution of 4 s for an entire volume of a cell, we devised a setup that allowed us to inject EGF or transferrin as a temporal pulse during image acquisition (Supplementary Fig. 1 and Supplementary Movie 1). 50 µL of 100 nM fluorescently labeled ligand was injected near the imaging region, followed by about 200-300 µL of imaging medium. Thus, in single acquisition sequences of up to 15 min, we could follow the initial binding of ligands to the receptors at the plasma membrane, the appearance of early endosomes formed directly by endocytosis of the ligands, and any of their further movements within the cell; additionally, APPL1 was tracked using APPL1-GFP. We confirmed that the EGF-bearing endosomes are not macropinosomes and are likely derived from clathrin mediated endocytosis (Supplementary Fig. 2). To independently track the coordinates of endosomes labeled with APPL1-eGFP, EGF or transferrin ligands, we performed 2-color (APPL1-GFP and Alexa 647 labeled transferrin or EGF) imaging of whole cells at 4 s per volume (Supplementary Movies 1,2). From the independently tracked coordinates of APPL1-GFP as compared to those of the two other ligands, we quantified the number of spatio-temporally correlated tracks (Methods, Supplementary Fig. 3). Both transferrin and EGF appeared as diffuse background fluorescence immediately following injection, and subsequently appeared as punctate structures on the plasma membrane (Fig. 1a,b). In the case of transferrin, the punctate structures slowly acquired low levels of APPL1, with an average delay of 100 s (Fig. 1b,g) (Supplementary Movie 2). In contrast, EGF immediately colocalized with APPL1 at the cell-periphery (Fig. 1a,d)), as indicated by the fraction of EGF tracks positive for APPL1 (Fig. 1d) and track towards the peri-nuclear region (Supplementary Movie 1).

**Fig. 1.**
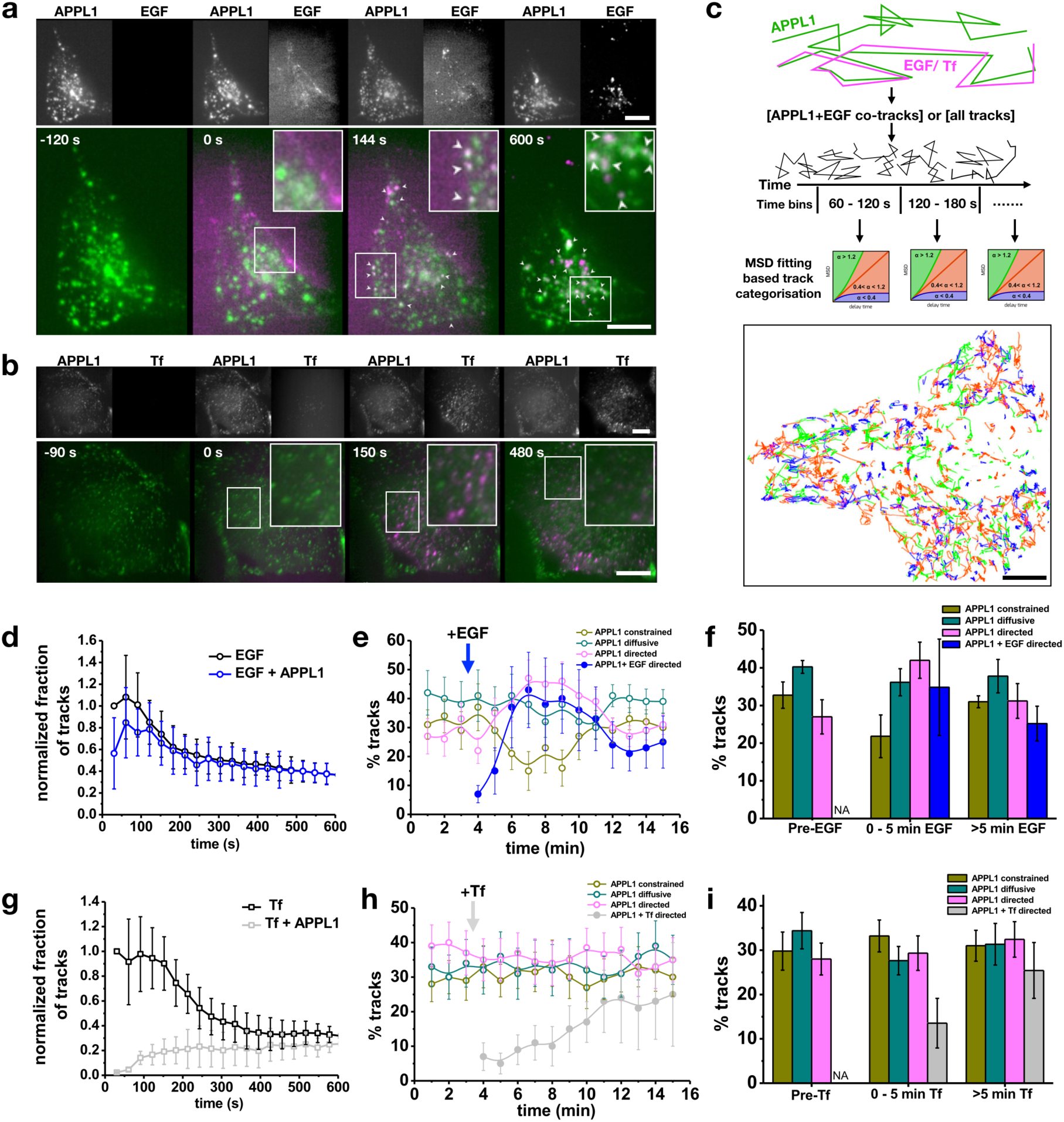
EGF rapidly colocalizes with APPL1 and is actively co-trafficked to the perinuclear region. **(a)** Cells transfected with APPL1-eGFP (green) were imaged using LLSM, during which time EGF-A647 (magenta) was pulse injected. The inserts correspond to APPL1 and EGFA647 from left to right for the denoted time points. Rectangles indicate zoomed versions for each time point. Time is in seconds. Arrows indicate APPL1 and EGFA647 colocalization. Scale bar = 10 µm. **(b)** Similar experiment with fluorescently labeled transferrin. No colocalization was observed. Scale bar = 15 µm. **(c)** Schematic of tracking and analysis workflow. All identified tracks, or tracks filtered on the basis of co-trafficking by presence of both channels within a determined radius sphere at each time point, were selected for ensemble MSD analysis. Based on the MSD analysis of the tracks, each track was characterized as constrained, diffusive or undergoing directed motion (Methods) and the tracks colored correspondingly (bottom) or exported for grouped statistical analysis. **(d)** Graph of fraction of cargo tracks with time (seconds) following EGF addition to HeLa cells transfected with APPL1 EGFP. Graphs show all the EGF tracks and the fraction of EGF tracks positive for APPL1. Error bars indicate standard deviation (n = 13 cells). **(e)** Percentages of APPL1 tracks categorized as constrained (yellow), diffusive (green) and directed (magenta) motions as a function of time by MSD analysis in a single cell. In addition, the blue trace shows the subset of APPL1 tracks positive for EGF that displayed directed motion, demonstrating that most APPL1 tracks with directed motions were also positive for the EGF. Error bars correspond to the standard deviation. **(f)** Percentage of APPL1 tracks undergoing constrained (yellow), diffusive (green), directed (magenta) motions and directed motions of EGF bearing APPL1 endosomes (blue) that were grouped in time as pre-cargo addition, 0-5 mins and 5-15 mins post addition based on example data presented in (e). Error bars correspond to the standard deviation where applicable (n = 9 cells). **(g)** Graph of the fraction of cargo tracks with time (seconds) following transferrin addition to HeLa cells transfected with APPL1 EGFP. Graphs show all the transferrin tracks and the fraction of transferrin tracks positive for APPL1. Error bars indicate standard deviation (n = 8 cells). **(h)** Percentages of APPL1 tracks categorized as constrained (yellow), diffusive (green) and directed (magenta) motions as a function of time by MSD analysis in a single cell. In addition, the graphs show the subset of APPL1 tracks positive for transferrin that display directed motion (grey), demonstrating that only a subset of APPL1 tracks showed the presence of transferrin in the initial time points post addition. Error bars correspond to the standard deviation. **(i)** Percentages of APPL1 tracks undergoing constrained (yellow), diffusive (green), directed (magenta) motions and directed motions of transferrin bearing APPL1 endosomes (grey) that are grouped in time as pre-cargo addition, 0-5 mins and 5-15 mins post addition based on example data presented in (e). Error bars correspond to the standard deviation where applicable (n = 7 cells).

To further analyze how APPL1 binding affects EGF bearing endosomes, we developed an analysis workflow that selected the APPL1 tracks that co-tracked with EGF (Fig. 1c). The overall motility characteristics of these tracks were then determined using mean square mean square displacement analysis [14]. In general, upon addition of EGF, the fraction of APPL1-EGF tracks exhibiting directed motion increased slightly while those with constrained motion decreased (Fig. 1e,f). However, by 5 min, of all APPL1 vesicles that showed directed motion, over 80% were also found to be positive for EGF (compare pink/blue curves, Fig. 1e, and columns, Fig. 1f), indicating that APPL1-EGF-bearing endosomes display a strong tendency towards being more mobile (Fig. 1 e,f). In contrast, by 5 min after the addition of transferrin, the APPL1-bearing endosomes most likely to show directed movements were those that lacked transferrin (Fig. 1h,i). EGF stimulation also resulted in APPL1-bearing endosomes exhibiting longer tracks, and higher maximum velocities, compared to unstimulated cells (Fig. 2a).

**Fig. 2.**
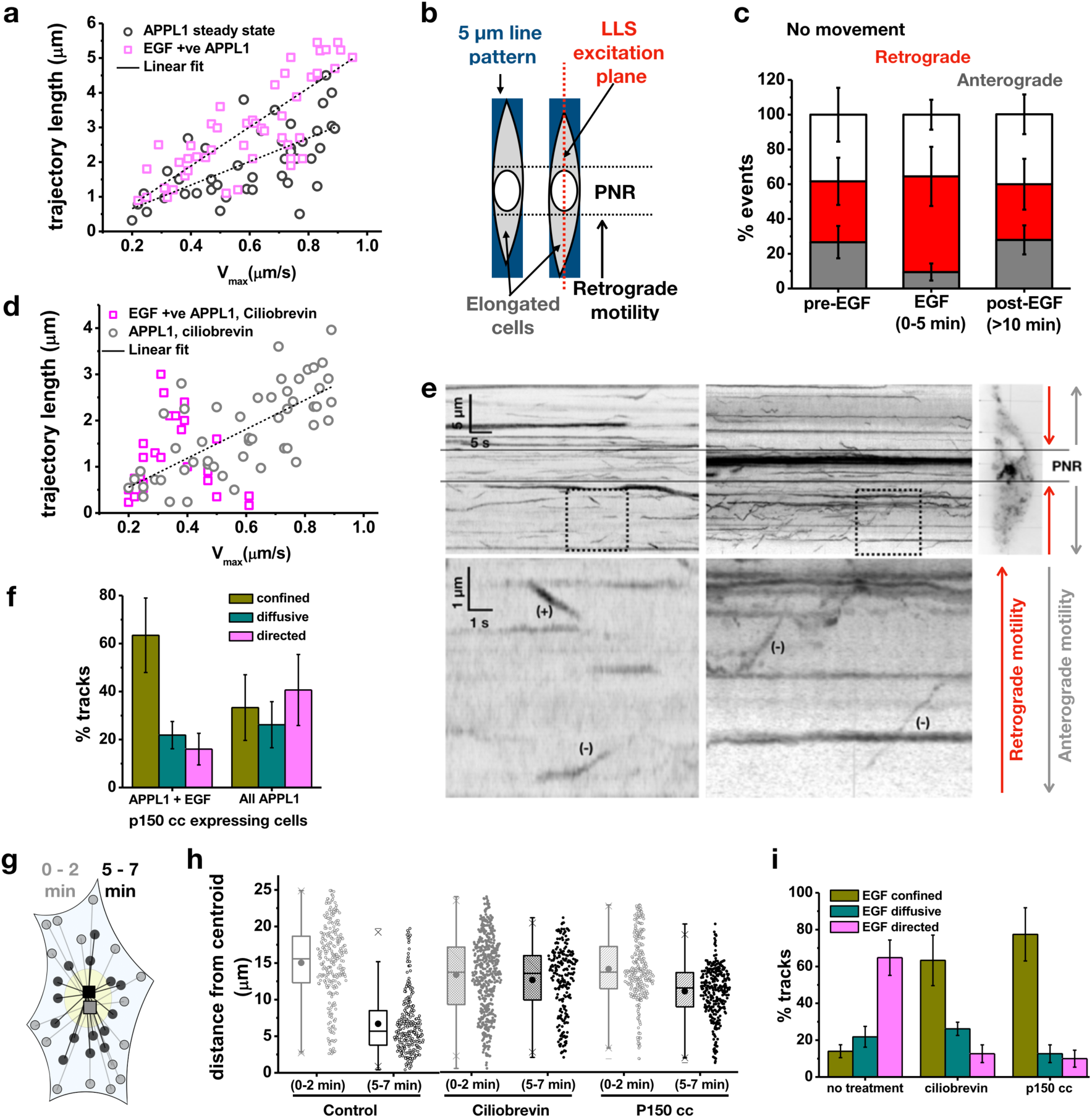
Dynein is responsible for rapid EGF-APPL1 trafficking to the perinuclear region. **(a)** Scatter plot of calculated trajectory lengths in microns against maximum velocity in um/s of APPL1 endosomes in unstimulated cells (black) and EGF-bearing APPL1 endosomes 2-7 mins following injection (magenta). Line of best fit plotted following linear regression. **(b)** Schematic of single plane lattice illumination (dotted red line) of cells micropatterned in 5 µm patterns (blue). This elongation of the cells accentuates retrograde motility towards the PNR as indicated by the arrow. **(c)** Percentages of APPL1 endosomes which underwent anterograde (grey), retrograde (red) or no net movement (white) in micropatterned HeLa cells pre-EGF stimulation, 0-5 mins and beyond 10 mins post-EGF stimulation. Error bars indicate standard deviation. **(d)** Scatter plots of the distance in microns of EGF endosomes from a selected centroid in HeLa cells, stimulated with 100 nM EGF, at 0-2 mins and 5-7 mins post addition. The grey plot represents the total amount of EGF bearing endosomes and the blue plots indicate APPL1 positive endosomes bearing EGF. The inner box of the box plot represents the standard deviation, the inner bar the median and the dot the mean, the ‘x’ represents the counts within 1-99% of the sample and the horizontal bars the range. **(e)** Kymographs of APPL1 endosomes in HeLa cells pre-EGF stimulation (left) and 0-5mins post-EGF stimulation (right). The top graphs show a zoomed-out view, scale bar = 5 µm y-axis, 5 seconds x-axis. The overlaid lines show the area of the kymograph corresponding to the perinuclear region as seen in the oblique slice insert. The bottom graphs show the area of the dotted box zoomed 5x, ‘+’ indicates plus-end directed motions ‘-’ indicates minus-end directed motions. **(f)** Percentages of EGF bearing endosome tracks which showed confined (yellow), diffusive (green) and directed (magenta) motion in HeLa cells 5-15 mins post stimulation with 100 nM EGF. The cells were either uninhibited, treated with 50 µM ciliobrevin or transfected with DsRed p150 cc. The error bars represent standard deviation. **(g)** Schematic of endosomal distribution calculation. Euclidian distances between endosomal positions (circles) and the centroid of all the positions (squares) were calculated within a time interval of 2 mins immediately post EGF stimulation (0 – 2 mins) and 5 mins later (5 – 7 mins). **(h)** Scatter plots of the endosomal distances from centroid in microns of EGF endosomes in HeLa cells stimulated with 100 nM EGF, at 0-2 mins and 5-7 mins post addition. The dotted box and whisker plots represent cells treated with 50 µM ciliobrevin and the stripped plots cells transfected with DsRed p150 cc. The inner box of the box plot represents the standard deviation, the inner bar the median and the dot the mean, the ‘x’ represents the counts within 1-99% of the sample and the horizontal bars the range. **(i)** A bar graph of the percentage of APPL1 (right) and APPL1 + EGF (left) bearing endosome tracks which show confined (yellow), diffusive (green) and directed (magenta) motion in HeLa cells treated with 50 µM ciliobrevin, 5-15 mins post stimulation with 100 nM EGF. The error bars represent standard deviation.

### EGF bearing APPL1 positive endosomes utilize dynein to move

To allow tracking at higher resolution we combined LLSM with micropatterning. On micropatterned coverslips with 5 µm lines, cells acquired an elongated shape (Methods and Supplementary Fig. 4). We predicted that the elongated shape of the cells would accentuate the directionality of endosomal movements, which, in the case of EGF-bearing APPL1 endosomes, are expected to occur in a PNR-directed, retrograde manner (Fig. 2b). We positioned the lattice light sheet such that an oblique cross-section illuminated the nucleus and adjacent PNR of each cell and acquired images with 200 ms of exposure per frame (Supplementary Movies 3, 4). The number of retrograde movements of APPL1 endosomes in wild-type cells treated with EGF was significantly higher than in unstimulated cells (Fig. 2c,e and Supplementary Movies 3, 4) indicating that EGF stimulation can cause a switch in APPL1 dynamics. While APPL1 is associated with endosomes that exist prior to EGF-stimulation, once it becomes attached to endosomes newly formed by the internalization of EGF, those endosomes are much more likely to move in a retrograde manner. These observations led us to ask three questions: What motors are engaged to enable retrograde movement? What causes APPL1 to switch from pre-existing endosomes to newly generated EGF-bearing endosomes? How does APPL1 localize to EGF bearing endosomes?

Given the observed retrograde motility of EGF-APPL1-bearing endosomes, we suspected that the main minus-end directed motor, dynein, might be involved. Dynein has also been implicated in the translocation of EGF to the juxta-nuclear region in earlier studies [28]. We expressed a fluorescently labeled version of p150 217 – 548 tagged with dsRed, which sequesters dynein, thus inhibiting dynein-dynactin-based motility [29]. This allowed us to select dsRed-labeled cells, and then assay the motility of their EGF and/or APPL1-bearing endosomes during the first 10 min post EGF addition (Supplementary Movie 5). Interestingly, while the total set of APPL1-bearing endosomes did not show a significant change in motility, in the subset that co-tracked with an EGF signal, motility was inhibited (Fig. 2f), with a majority of the tracks showing constrained motion (>60%) and only about 15 % showing directed motion (Fig. 2f). We also quantified the extent of accumulation of endosomes at the PNR by measuring the distance from the centroid, for all endosomal populations (Methods and Fig. 2g). In contrast to controls, no accumulation of EGF vesicles in the PNR was observed when cells expressed p150 217 – 548 or were treated with another dynein inhibitor, ciliobrevin (Fig. 2h). These results indicate that dynein is the major motor protein involved in the translocation of these endosomes. In agreement with this, the EGF-bearing endosomes exhibited a significantly increased fraction of constrained movements under dynein inhibition, in contrast to the large fraction of directed movements seen in untreated cells (Fig. 2i). Therefore, independent experiments, that perturb dynein in distinct ways, both suggest that the accumulation of EGF-APPL1-bearing endosomes in the PNR, via minus-end directed translocation, requires the recruitment of dynein.

### APPL1 mediated translocation to the perinuclear region is necessary for efficient maturation of EGF bearing endosomes

Maturation of APPL1 positive endosomes have been shown to occur through a process of conversion, where one protein is shed off the surface of the endosome, and a new endosomal marker is acquired. APPL1 endosomes convert into EEA1 positive endosomes. This has also been demonstrated for EGFR containing endosomes. We first recapitulated the maturation of EGF bearing endosomes (Fig. 3a-d). With time, the number of EGF tracks that acquired EEA1 increased with time. Within 5 minutes, a steady fraction of EGF-EEA1 double positive endosome is achieved (Fig. 3d). As previously described, conversion processes where APPL1 is shed off and EEA1 is acquired were also observed (Fig. 3c). Inhibiting retrograde motility by treatment of cells with Ciliobrevin slowed down APPL1 to EEA1 maturation of EGF bearing endosomes (Fig. 3d). The localization of EGF-APPL1 double positive endosomes therefore, is necessary to maintain the flux of EGF bearing endosomes into EGF-EEA1 double positive endosomes through conversion.

**Fig. 3.**
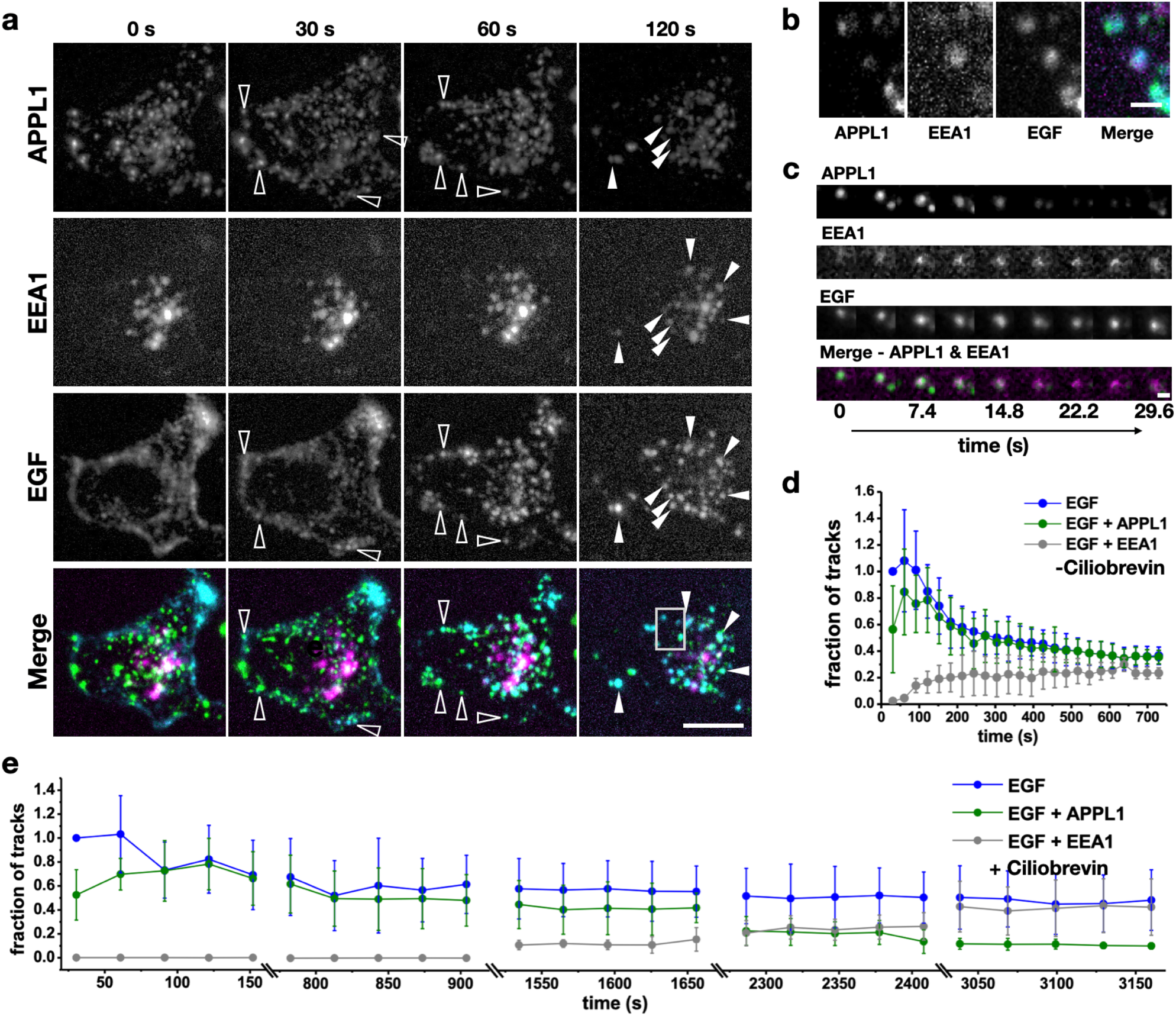
APPL1-EGF endosomes mature by acquiring EEA1 and is dependent on its retrograde motility and localization to PNR. **(a)** Representative maximum intensity projections of 100 mM EGF stimulation of HeLa cells expressing APPL1-eGFP and EEA1 TagRFP-T (Magenta). Hollow arrows point at few examples of APPL1 and EGF colocalization and the solid arrows point at APPL1-EEA-EGF1 triple or EEA1-EGF double positive endosomes. Scale bar = 15 µm **(b)** A zoomed view of APPL1-EEA1-EGF triple positive endosomes in the boxed region in (a). Scale bar = 0.5 µm **(c)** An EGF-APPL1 double positive endosome undergoing maturation by conversion to EGF-EEA1 positive endosome. Scale bar = 0.2 µm **(d)** Graph of fraction EGF tracks positive for APPL1 and EEA1. Error bars indicate standard deviation (n = 6 cells). **(e)** Same plot as (d) for cells treated with 50 µM Ciliobrevin. Double lines on the X-axis indicate breaks in time.

### APPL1 desorption is dependent on calcium wave evoked by EGF stimulation

To investigate the observed recruitment of APPL1, from pre-existing to newly formed EGF-bearing endosomes, we analyzed the whole-cell dynamics of APPL1. Upon stimulation with 100 nM EGF, the APPL1 signal was lost from the endosomes of the PNR (Fig. 4a,b and Supplementary Movie 6, 7) suggesting global redistribution of APPL1. We used Imaris to segment out the PNR to quantify the loss of APPL1 signal from the endosomes. We chose the endosomes at the PNR for this measurement as they are relatively immobile within the time frame of APPL1 desorption, as compared to the peripheral endosomes (Supplementary Movie 8). We found that PNR endosomes lose their APPL1 signal completely or partially when stimulated, respectively, with 100 nM EGF or 20 nM EGF (Fig. 4c). In contrast, upon transferrin stimulation, we did not observe any APPL1 redistribution. EGF stimulation has been shown to elicit Ca^2+^ signals through phospholipase cγ (PLCγ) and phospholipase A2 (PLA2) [12, 13]. We speculated that the transient increase in cytosolic Ca^2+^ evoked by EGF binding could cause the loss of APPL1 from the endosomes. Previous studies of other proteins that, like APPL1, contain a pleckstrin homology (PH) domain, have shown that elevated intracellular Ca^2+^ prevented membrane binding of such proteins through the formation of Ca-phosphoinositide complexes [30]. To determine if elevated Ca^2+^ is responsible for the unbinding of APPL1, we imaged the dynamic distribution of APPL1 GFP and the Ca^2+^indicator R-GECO, after using 100 nM ionomycin to induce a Ca^2+^ influx (Fig. 4d and Supplementary Movie 9) [31]. Upon ionomycin addition, APPL1 dissociated from endosomes within 40 s of the R-GECO signal peak (Fig. 4d). In all cases of ionomycin-mediated desorption, we observed that the endosomes became transiently immobile, after which APPL1 desorbed from them before subsequently rebinding to endosomes. We attribute the freezing of motile endosomes upon an increase in cytosolic Ca^2+^ to ‘Calcium-mediated Actin Reset’ (CaAR), which results in a dense meshwork of actin [32]. We also measured the time of R-GECO peaks with respect to EGF binding, and APPL1 desorption with respect to EGF binding (Fig. 4e and Supplementary Movie 10). We found the R-GECO signal peaked about 30 s post EGF stimulation, and APPL1 desorption occurred in the next 30 s (Fig. 4e).

**Fig. 4.**
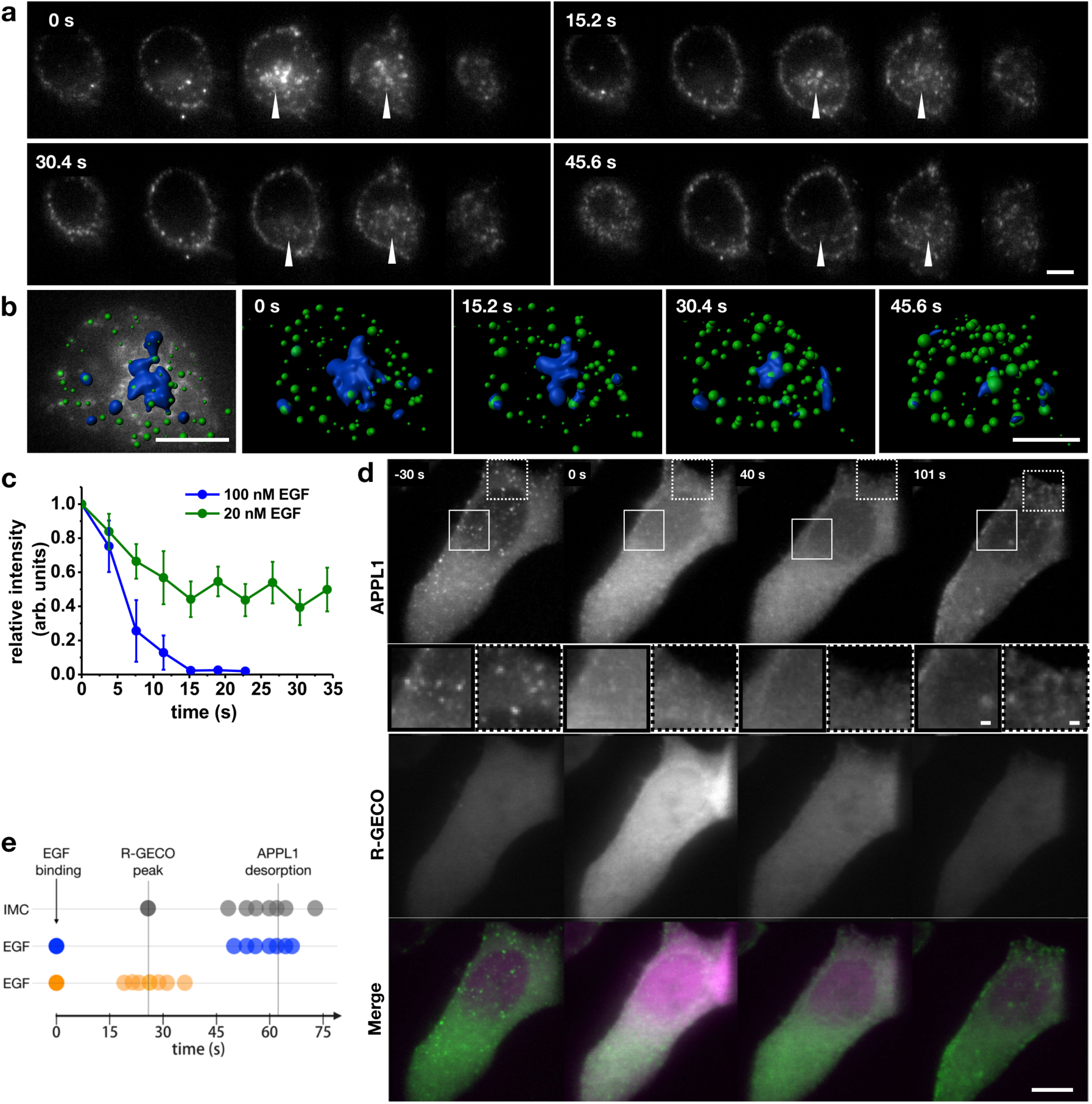
EGF mediated Ca^2+^ influx causes transient APPL1 desorption. **(a)** Time-lapse montage of cross-sections across a single cell displaying desorption of APPL1-eGFP following addition of 100 nM EGF. Scale bar = 4 µm. **(b)** Example image depicting segmentation of APPL1 eGFP signal in the PNR using Imaris and its loss upon EGF stimulation. Scale bar: 5 µm. **(c)** Change in relative intensity of APPL1-GFP in the perinuclear region of HeLa cells following addition of an EGF pulse of either 100 nM (blue) or 20 nM (green). Time is recorded in seconds following EGF binding, error bars represent standard deviation. **(d)** Maximum intensity projection of images of APPL1-eGFP (top, green in Merge) and R-GECO (middle, magenta in Merge) in HeLa cells stimulated with 100 nM pulse of ionomycin. Time recorded in seconds from ionomycin addition. Scale bar = 10 µm. Inserts correspond to zoomed sections (unbroken and dotted squares respectively) at each time point, scale bar = 1 µm. **(e)** Timeline of relationships between EGF binding, R-GECO peaks and APPL1 desorption. Measured R-GECO fluorescence peak and APPL1 desorption (grey dots), each dot represents a single experiment, with R-GECO peaks overlaid to mean of EGF induced R-GECO peaks. 100 nM EGF was used to determine both the time of APPL1 desorption (blue) and R-GECO peak (orange).

### APPL1 binds to EGFR through its PTB domain

It has been previously reported that APPL1 can bind directly to phosphorylated EGFR through its PTB domain. To test whether this interaction plays a role in early stage trafficking of EGFR, cells expressing WT-APPL1 were treated with 10 µM erlotinib for 1 h, to inhibit EGFR cross-phosphorylation [33], and imaged following EGF stimulation (Fig. 5a). APPL1 localization in these cells was extremely rare and transient. To investigate whether the PTB domain of APPL1 is specifically involved in the binding to EGFR, as suggested elsewhere [25], we expressed a mutant of APPL1 with the PTB domain deleted (APPL1-ΔPTB) [34]. We found that APPL1-ΔPTB did not localize to EGF-bearing endosomes and the signals were cytoplasmic. Furthermore, in both experiments, in cells either expressing APPL1-ΔPTB or treated with erlotinib, the peri-nuclear localization and directed motion of EGF-bearing endosomes were strikingly impaired, as demonstrated by the abrogation of the decrease in mean centroid to endosomal distances (Fig. 5 b,c). Some mobile endosomes were observed, possibly attributable to kinesins involved in EGFR trafficking [35]. In the case of cells expressing APPL1-ΔPTB, the impaired localization of EGF endosomes in the PNR strongly suggests that this deletion mutant functions as a dominant negative. The experiments together suggest that APPL1 is an intermediary between phosphorylated EGFRs and dynein and is essential for perinuclear accumulation.

**Fig. 5.**
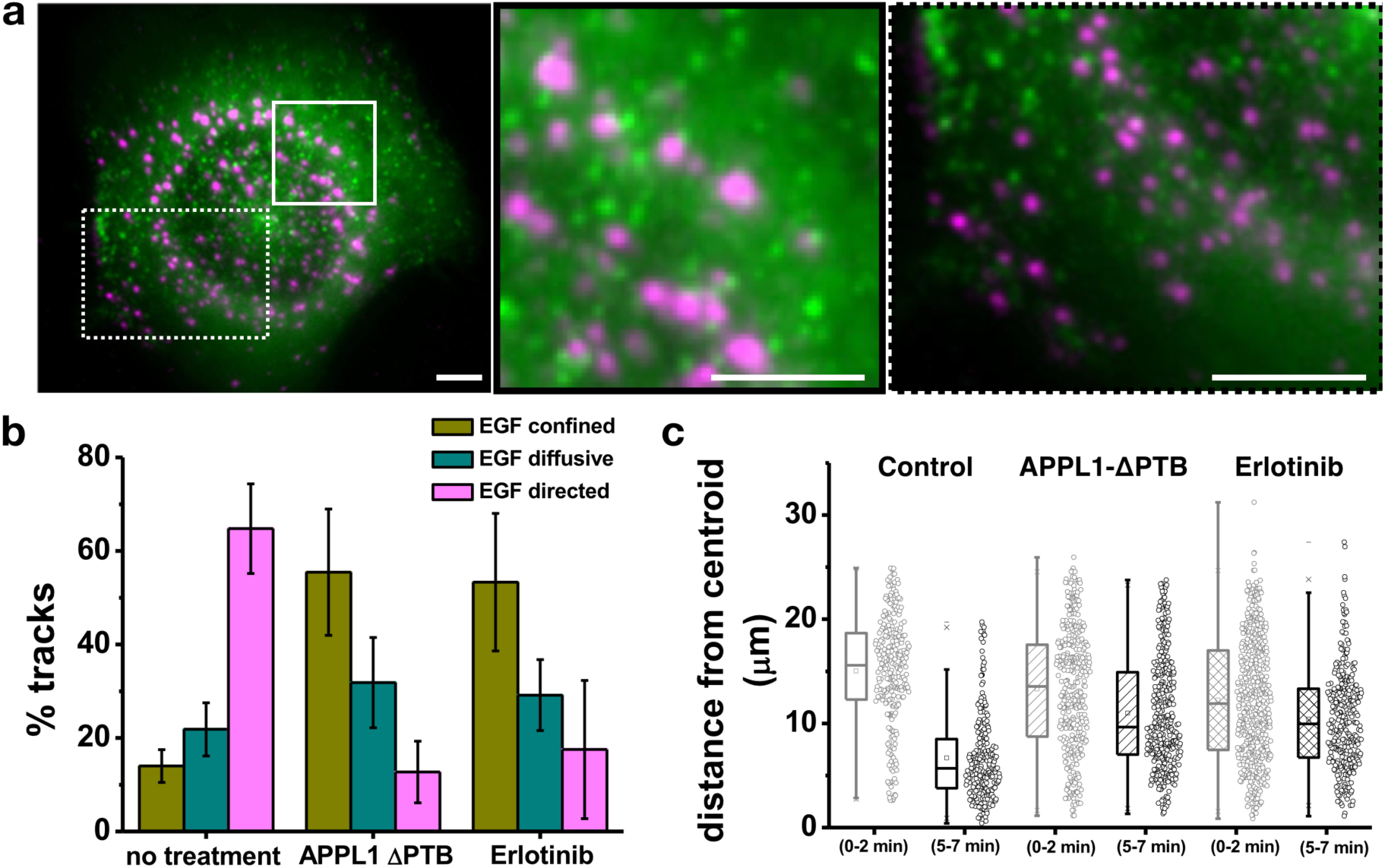
APPL1 binds activated EGFR via its PTB domain to mediate minus-directed motility. **(a)** Representative maximum intensity projection of LLSM imaging of WT-APPL1-GFP (green) in HeLa cells treated with erlotinib 5 min post 100 nM EGF-647 (magenta) stimulation. Scale bar = 4 µm. Left and right panels indicate zoomed sections corresponding to unbroken and dotted rectangles respectively, scale bar = 4 µm. **(b)** Percentage of EGF bearing endosome tracks which showed confined (yellow), diffusive (green) or directed (magenta) motility in HeLa cells transfected with APPL1-eGFP (no treatment), APPL1-ΔPTB eGFP or APPL1-eGFP and treated with 10 µM erlotinib. Error bars correspond to the standard deviation. **(c)** Scatter plots of EGF distance in microns from centroid in HeLa cells treated with 100 nM EGF at 0-2 mins and 5-7-mins post addition as described in Fig. 2g. The Hela cells were either transfected with APPL1-GFP (blank plot), APPL1-ΔPTB GFP (diagonal lined plot) or APPL1-GFP and treated with 10 µM erlotinib (crossed plot). The inner box of the box plot represents the standard deviation, the inner bar the median and the dot the mean, the ‘x’ represents the counts within 1-99% of the sample and the horizontal bars the range.

## DISCUSSION

### A mechanism for cohort transport of signaling EGFRs

Based on the aforementioned experiments that provide evidence of endosomal dynamics and protein redistributions, in Fig. 6 we present a model of the proposed events involving EGF, EGFR, Ca^2+^, APPL1, and dynein from before, and up to 5 min after, EGF stimulation. Before exposure to EGF, EGFR exists in its monomeric state [36], APPL1 is bound to pre-existing endosomes, and Ca^2+^ levels are low. Within 30 s of EGF-EGFR binding, EGFR dimerizes and is phosphorylated, endocytosis generates the first EGF-bearing vesicles, and Ca^2+^ levels begin to rise. Rising Ca^2+^ levels result in the desorption of APPL1 from pre-existing endosomes presumably by blocking the interaction of the BAR and PH domains of APPL1 and endosomal phosphoinositides, a type of block reported for other PH-domain containing proteins, including Akt [30]. Extracellular EGF concentration controls the degree of Ca^2+^ elevation [13] and in turn, the extent of APPL1 desorption, thereby resulting in a transient availability of cytosolic APPL1 proportional to the EGF concentration as demonstrated by the dependence of APPL1 desorption on EGF concentration. The resulting transient increase in cytosolic APPL1 occurs within approximately 60 s post EGF-binding, within which time EGFR phosphorylation continues in parallel. Previously, EGFR phosphorylation dynamics, measured using a ratio-metric sensor based on EGFR – ECFP and PTB-YFP, revealed that phosphorylation occurred in less than a minute after EGF addition [37]. This is consistent with the dynamics of APPL1 localization to EGFR bearing endosomes, through the PTB domain, observed in our experiments.

**Fig. 6.**
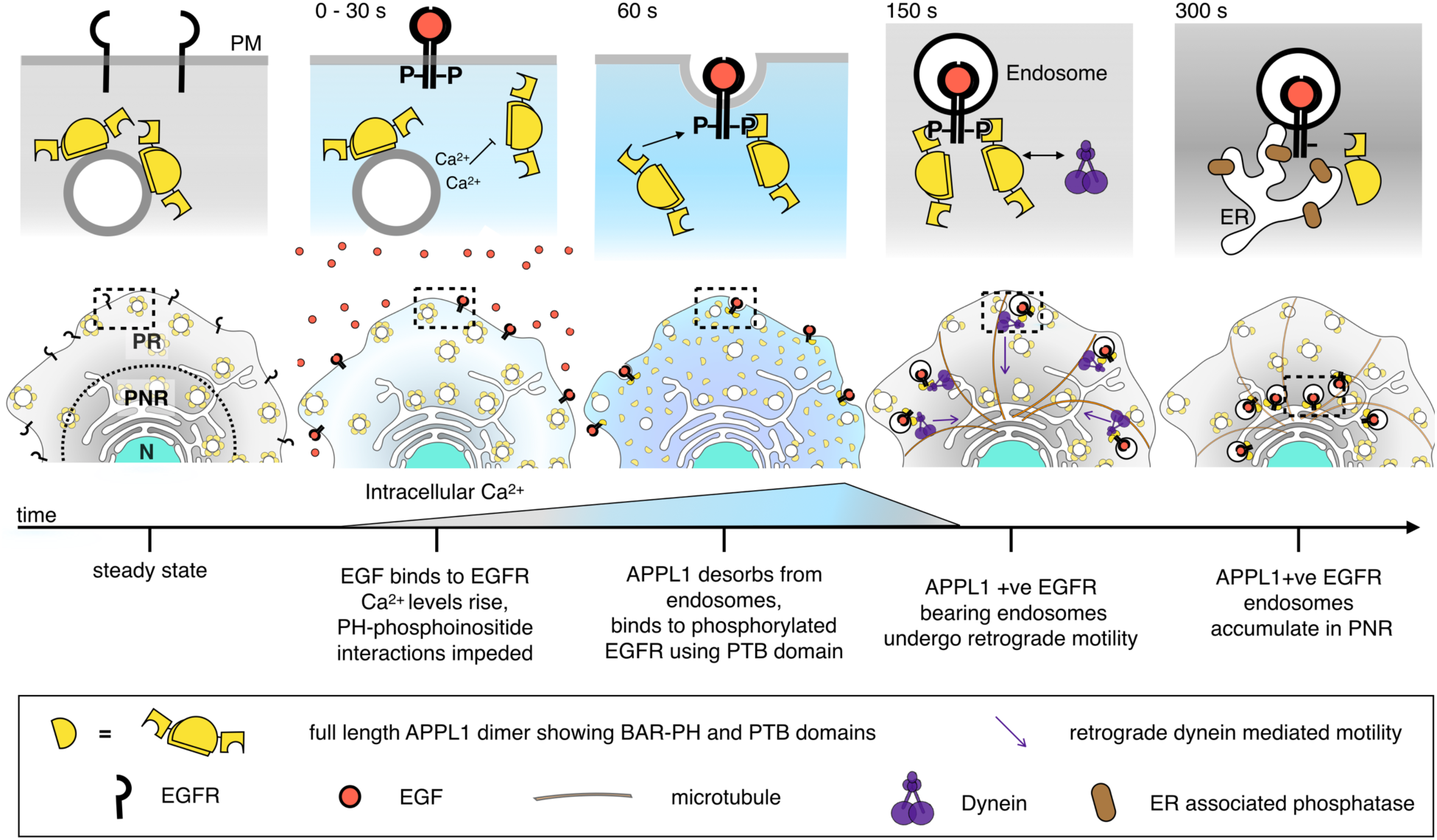
Summary figure of proposed mechanism of APPL1-mediated shunt trafficking of activated EGFR. In steady state cells, APPL1 (yellow semi-circles) is dispersed throughout the cell. EGF binding leads to dimerization and phosphorylation of EGFR and an increase in intracellular Ca^2+^ (blue background). The increase in intracellular Ca^2+^ impairs APPL1 PH binding to phosphoinositide causing APPL1 desorption. The increased cytosolic APPL1 results in APPL1 binding to activated EGFR via a PTB domain. APPL1 positive EGF bearing endosomes undergo dynein (purple figure) mediated motility to the perinuclear region which is rich in ER.

### APPL1 as a multifunctional endosomal signaling adaptor protein

The multiple interaction sites provided by APPL1’s BAR, PH and PTB domains are central to the model. The PTB domain of APPL1 is at the C-terminus and is structurally similar to the PTB domain of Shc [22], which also binds EGFR [38]. The PTB domain of Shc recognizes NPX(phosphor)Y and has been found to bind tighter to phosphorylated peptides than to phosphatidylinositol [39] and may explain the preference of APPL1 binding to phosphorylated EGFR than the membranes though phosphoinositides. Our experiments with APPL1-ΔPTB, where APPL1 binding to EGF bearing endosomes was impaired, support the fact that the PTB domain of APPL1 binds directly to phosphorylated EGFR. Once APPL1 has bound to EGFR, a form of yet unknown interaction appears to engage dynein, and EGF-bearing endosomes are rapidly shunted to the PNR. Our results suggest that dynein acts downstream to APPL1, in that its activity occurs after APPL1 recruitment, and when APPL1 was blocked from binding to EGFR (by erlotinib or APPL1-ΔPTB), no retrograde motility was seen. Dynein has previously been reported to be required for the translocation of EGF towards the nucleus [28]. While direct interactions between dynein and APPL1 have not been reported, thirteen key phosphorylation sites have been identified on APPL1, and their functions are yet to be fully elucidated [40]. How APPL1 coupled to EGFR promotes dynein engagement is unknown, however it is possible that APPL1 phosphorylation sites are involved. APPL1 phosphorylation has previously been implicated in regulating the recycling of activated GPCRs [41]. Collectively these studies highlight the importance of APPL1 phosphorylation and the need for further investigation of the multiple roles of APPL1 in regulating endosomes.

### Endosomal trafficking and signaling

What could be the physiological significance of such rapid trafficking of the EGFR receptors? Endocytosis and trafficking of signaling receptors regulate signal transduction by providing specificity in space and time [3, 36]. Endosomal trafficking allows signaling receptors to be localized at specific regions of the cell or to be channeled, in a timed fashion, towards attenuation sites in multi-vesicular bodies or lysosomes [6]. A recent report by demonstrates that EGFR signaling is controlled by protein tyrosine phosphatases (PTPs) associated with ER, that are enriched in the PNR [8]. The ER associated PTPs (PTP1B and TCPTP) enable spatially restricted activity and efficiently de-phosphorylate the EGFR when EGFR localizes to the PNR [4, 8]. Our results also agree with the distinct rapid motility previously reported for EGF in contrast to transferrin [27], and reveal the APPL1 mediated engagement of dynein as a mechanism for these movements. This study reveals that APPL1 is central to this dynein dependent rapid transport, as well as to the peri-nuclear accumulation of stimulated EGFRs and provides a mechanism distinct from that used by constitutive receptors like transferrin. A recent study also reported APPL1 displaying fast retrograde movement in axonal transport hinting that the dynein-based motility that we describe may be crucial in different cell types in different contexts [42].

Vesicular transport is an important logistical support in the signal processing machinery of the cell. As mentioned above, localization of receptors at the PNR, that results in dephosphorylation and hence signal attenuation is an important step in the signal processing. It is likely that not just the localization, but the rate of localization from the periphery of the cell to the PNR will determine the signal sensing. In a scenario where the receptors depend on the steady state dynamics of the endosomal dynamics, the arrival times of the receptors at the PNR may be broadly distributed. However, as described in this study, if the receptor depends on a specific activated mechanism where a cohort movement of receptors ensue, the arrival times may be restricted to a narrow distribution.

## CONCLUSION

Our findings also highlight the sophisticated organization of the endosomal system’s interaction matrix, where transient interactions driven by specific biochemistry take place in timescales of minutes. Our approach reveals a previously unknown EGFR-APPL1-dynein nexus and highlights how live cell imaging-based studies can unravel transient, conditional interactions, capturing simultaneous multiple processes triggered by a single protein. Such transient interactions are perhaps not discernible in ensemble approaches where temporal resolution, and details of endosomal specificity and dynamics, are lost.

## Methods

### Cell lines

HeLa and RPE1 (ATCC) cells were incubated at 37°C in 5% CO_2_ in high glucose Dulbecco’s modified eagle media (DMEM) (Life Technologies), supplemented with 10% fetal bovine serum (FBS) and 1% penicillin and streptomycin (Life Technologies). Cells were seeded at a density of 200,000 per well in a 6-well plate containing 5 mm glass coverslips.

### Live-cell imaging

Cells were imaged using a lattice light sheet microscope (3i, Denver, Colarado, USA). Excitation was achieved using 488, 560 and 640 nm diode lasers MPB communications) at 1-5% AOTF transmittance through an excitation objective (Special Optics 28.6× 0.7 NA 3.74-mm immersion lens) and is detected via a Nikon CFI Apo LWD 25x 1.1 NA water immersion lens with a 2.5x tube lens. Live cells were imaged in 8 mL of 37°C-heated DMEM and were acquired with 2x Hamamatsu Orca Flash 4.0 V2 sCMOS cameras using Latticescope. 600 µL of 100 nM Alexa647 labelled EGF or transferrin, or 600 µL of 100 µg/µL dextran-fluorescein was added mid imaging using a custom syringe-sample holder contraption that allowed precise injection of the desired volume of fluorescently labelled ligands, followed by imaging media. The fluorescently labelled ligand containing media were kept separated from the imaging media by an air bubble (Supplementary Fig. 1). Similarly, non-fluorescent 600 µL of 100 nM ionomycin was added where indicated. Cells were serum starved in high glucose DMEM medium 4 h prior to EGF addition.

### Plasmids and transfection

Cells were transfected with pEGFPC1-human APPL1, a gift from Pietro De Camilli (Addgene plasmid #22198) (26), GFP-APPL1-ΔPTB, a gift from Donna Webb (Addgene plasmid #59768) (34), DsRed-p150 217-548, a gift from Trina Schroer (Addgene plasmid #51146) (29), EEA1 TagRFP-T (Addgene plasmid #42635), a gift from Silvia Corvera. Cells were transfected with a total of 1 µg DNA (plasmid of interest – 0.2 µg, blank DNA – 0.8 µg) for single protein expression or (plasmids of interest – 0.2 µg + 0.2 µg, blank DNA – 0.6 µg) using lipofectamine 3000 (Thermo Fisher Scientific) or the Neon electroporator (Invitrogen) at 1200 mV 20 ms, 2 pulses for RPE1 and 1005 mV 35 ms, 2 pulses for HeLa. HeLa cells were ensured for mild expression by the following: The transfection mix consisted of APPL1 or EEA1 plasmids adding up to 20 % of the total DNA amount for transfection, with the rest consisting of blank DNA. Secondly, it has been reported that overexpression of APPL1 or EEA1 results in colocalization of APPL1 and EEA1 on Rab5 endosomes. We optimized this concentration by screening for this artifact, where we see no overlap of APPL1 and EEA1. Thirdly, APPL1 overexpression impairs EGFR internalization. However, in all our experiments, EGF/EGFR complex is trafficked efficiently to the peri-nuclear region.

### Micropatterning

Coverslips were sonicated with 70% ethanol for 30 min and plasma cleaned for 5 min. Coverslips were incubated with 1 mg/mL poly(L-lysine)-poly(ethylene-glycol) (PLL-g-PEG) in 1x phosphate buffered saline (PBS) at 4°C overnight. Coverslips were then placed on a chromium mask with 5 µm line patterns and illuminated for 5 min by an ultraviolet lamp. The coverslips were then incubated with 20 µg/mL fibronectin (Thermo Fisher Scientific) in 1x PBS with 0.02% Tween 20 and 0.04% glycerol for 1 h at room temperature. Following which cells were plated onto the slips and allowed to take the shape desired over 4 h (Supplementary Fig. 4).

### Drug addition

Cells were incubated with ciliobrevin (CB) or dimethyl sulfoxide (DMSO) at a final concentration of 200 nM in 8 mL DMEM medium for 5 min before and during imaging. Cells were similarly treated with 100 nM final concentration of phorbol12-myristate-13-acetate (PMA) or PBS; and 10 µM erlotinib or DMSO control for 1 h prior to imaging as indicated.

### Co-tracking analysis

Imaris 9.2.1 (Bitplane) was used to detect and track vesicles via the spot detection feature following Gaussian filtering 0.1 um. The track coordinates were exported and analyzed using custom codes developed in-house. Tracks which followed the same trajectory were filtered by the following routine: The errors in the measurement between the two channels as acquired by the two cameras were obtained by diffusing Tetraspeck beads (T7279, Thermo Fischer Scientific). This estimation of error provided the minimum radius for analyzing the ‘co-localization’ of spots belonging to two distinct trajectories in two different channels. The spots were considered colocalizing if they were within a sphere defined by the minimum radius: and a time filter of 10 consecutive frames, where p = x, y, z coordinates as a function of time for one channel and c for another (Supplementary Fig. 3). The effective radius of co-localization was set to be 500 nm to account for the sequential imaging and any spatial segregation within a single endosome, based on the measurements from diffusing beads. Next, once all the colocalizing spots were identified, using the trajectory ID of the spots, they were filtered using a minimum time window of 3 frames (12 s). We discarded any tracks that co-tracked for less than 12 s as they could be due to transient interactions. Tables detailing the number of co-tracks identified with time were exported, averaged for many samples and plotted using ORIGIN.

### Motility analysis

The track coordinates were exported from Imaris. MSD analysis was performed as described in previous studies [43, 44]. Briefly, the identified trajectories were used to generate MSD plots. The MSD curves were fitted with the following equation: *MSD* = < *R* >^2^= 4*Dt*^α^ + 2σ^2^, where D is the diffusion coefficient, t is the lag-time, α, the scaling exponent, and σ is the measurement error measured on the LLSM using beads adsorbed onto glass, fixed as 35 nm for the fits. The trajectories were characterized according to the scaling exponent α, retrieved from the fits. Tracks with an exponential value of <0.4 were termed as constrained, those with an exponential value of 0.4< α <1.2 termed diffusive and those with a value >1.2 were said to undergo directed motion.

### Peri-nuclear localization (distance from centroid)

Endosomal distributions within the cell were quantified by calculating the centroid of all endosomal localizations for a given time point. The Euclidian distances of each endosome from this centroid were calculated. These centroid-to-endosome distances were pooled, and ensemble statistical analysis was performed. Peri-nuclear accumulation resulted in lower distances between the centroid and each endosomal localization.

## Author contribution

SA devised and directed the study, HY, AK, SA performed LLSM experiments. HY, AP, SA analyzed data. AB performed the micropatterning, AK, UK, HG contributed to molecular biology, SA, HY and GH wrote the manuscript.

## Acknowledgements

The authors would like to thank Profs. Joe Howard, Marino Zerial for engaging discussions and Drs. Vaishnavi Ananthanarayanan, Angika Basant and Robert Weatheritt for valuable comments on the manuscript. This work was supported by ARC LIEF, LE150100163 awarded to Prof. Katharina Gaus.

