## Supplementary Material for "Rapid Whole Cell Imaging Reveals A Calcium-APPL1-Dynein Nexus That Regulates Cohort Trafficking of Stimulated EGF Receptors"

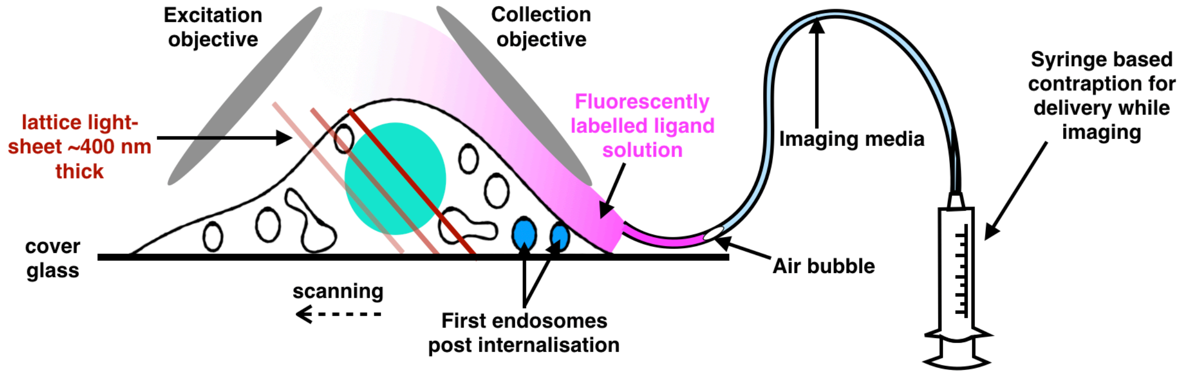


**Supplementary figure 1. Cartoon schematic of imaging and cargo delivery arrangement**

Cells are excited by a ~400 nm thick multi-Bessel based light sheet and imaged by a perpendicular collection objective during piezo scanning to visualize the entire cell. Cargo is added via a syringe contraption which has an outlet adjacent to the coverslip allowing for direct addition during image acquisition. Given that the volume of the LLSM's sample chamber is 9 ml, sufficient dilution of the injected media was achieved within 2 minutes (supplementary movie 1).

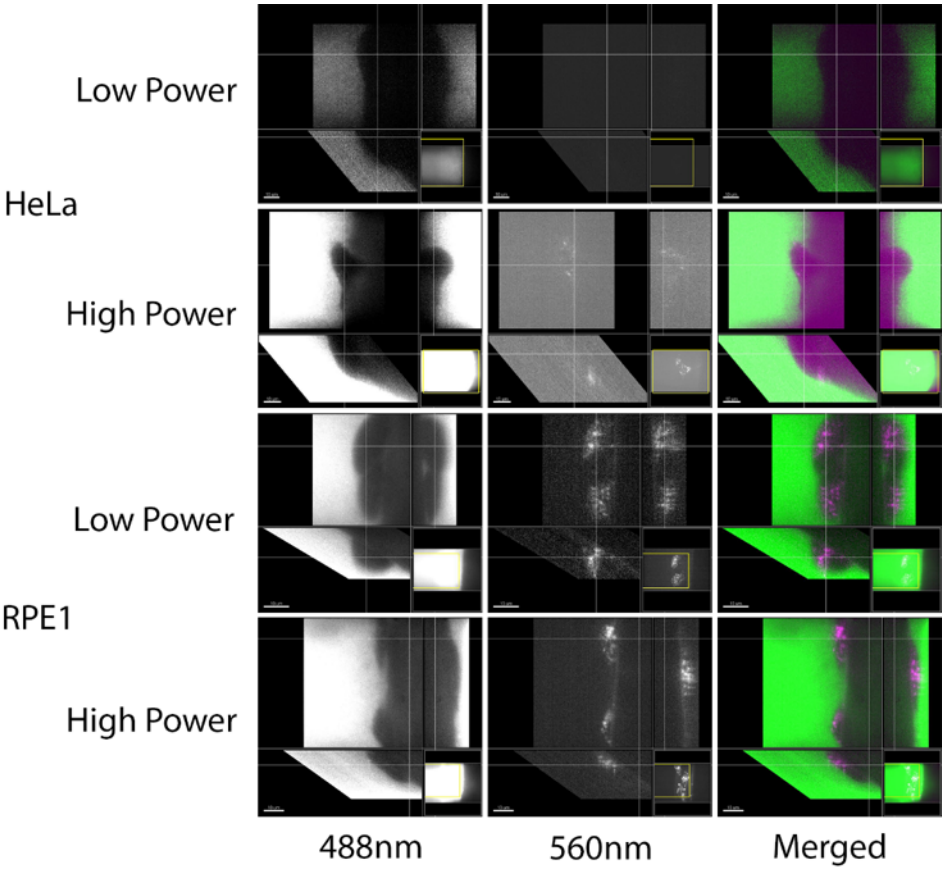


**Summary figure 2. HeLa cells do not form macropinosomes.**
Representative orthogonal views of LLSM imaging of HeLa (top) and RPE1 (bottom) cells following 100 µg/µL Dextran-fluorescein injection under both low (1%) and high (25%) laser power as indicated. The 488nm channel indicates non-internalized dextran while the 560nm channel indicates internalized dextran as fluorescein is a ratiometric probe for pH that reports on the alkaline endosomes when internalized [1].

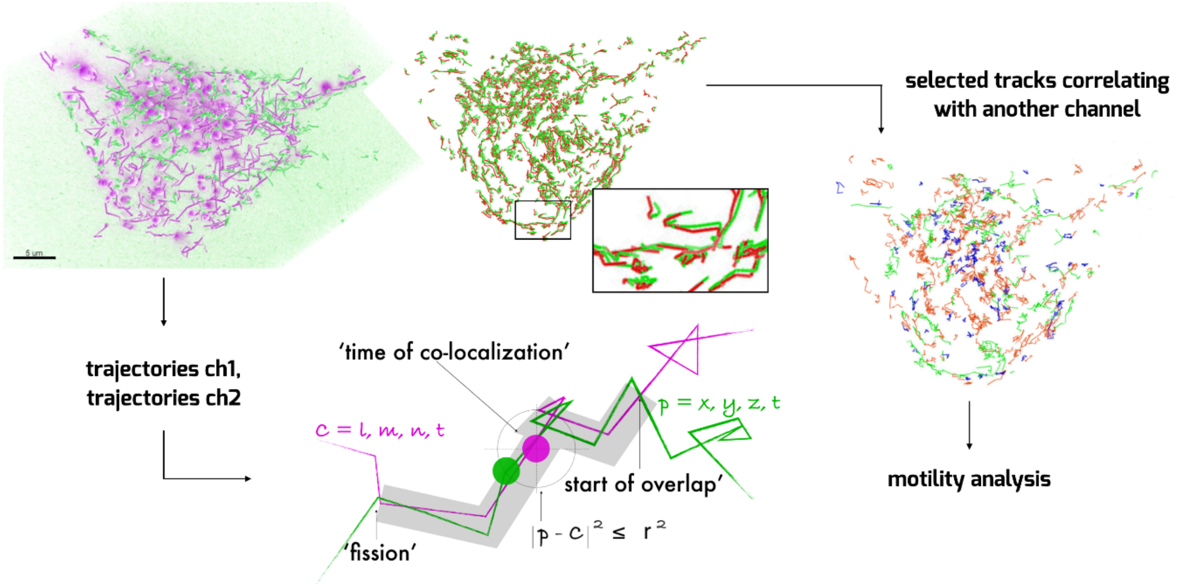


**Summary figure 3. Co-tracking analysis**

Image of detected tracks for two channels (green and magenta). Co-tracking was analyzed by setting the following condition between tracks: $\left|(\mathrm{p}-\mathrm{c})\right|^{2} \leq r^{2}$ and a time filter of 10 consecutive frames, where p = x, y, z coordinates as a function of time for one channel and c for another. The effective radius of co-localization sphere was set to be 500 nm to account for the sequential imaging and any spatial segregation within a single endosome. The identified tracks that co-track with the second channel can be extracted for further analysis of conditional motility, i.e. motility analysis of tracks showing co-localization of a second channel (Supplementary Movie 3).

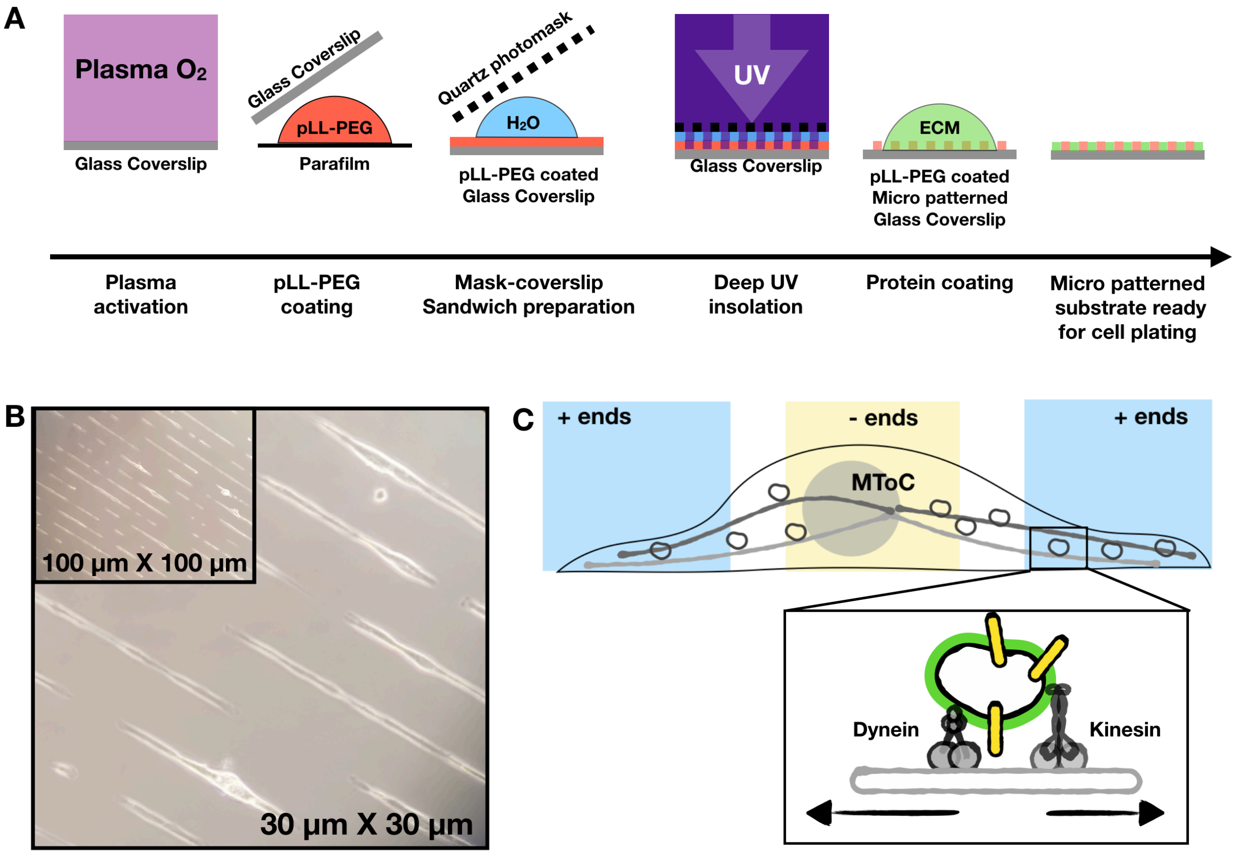


**Supplementary figure 4. Micropatterning of coverslips.**
(A) Schematic representation of micropatterning process of coverslip (based on [2]). (B) Example image of HeLa cells micropatterned on 5 µm lines, imaged using bright field microscopy at 100x and 30x (insert) magnification. (C) Schematic of microtubule arrangement in an elongated cell highlighting the peripheral plus ends (blue) and peri-nuclear minus ends (yellow) near the microtubule organizing center. Micropatterning cells into thin lines leads to alignment of microtubules within a cell. Rectangular insert corresponds to an APPL1 positive endosome with expected directionality for dynein-based motility and kinesin-based motility with respect to the elongated shape (Supplementary Movie 4).

### Supplementary information.

**Legend Movie 1**
Lattice light-sheet imaging of HeLa transfected with APPL1 EGFP and Alexa 647 labelled EGF addition while imaging.

**Legend Movie 2**
Lattice light-sheet imaging of HeLa transfected with APPL1 EGFP and Alexa 647 labelled Tf addition while imaging.

**Legend Movie 3**
APPL1 movements in unstimulated cells, single oblique cross section.

**Legend Movie 4**
APPL1 movements in EGF stimulated cells, single oblique cross section.

**Legend Movie 5**
APPL1 motility and EGF motility in cells expressing p150 cc dsRed.

**Legend Movie 6**
Oblique cross sections of a single cell demonstrating loss of APPL1 signal at PNR upon EGF stimulation.

**Legend Movie 7**
Two cells demonstrating whole-cell redistribution with loss of APPL1 signal at PNR upon EGF stimulation.

**Legend Movie 8**
Segmentation of the PNR to quantify loss of APPL1 signal upon EGF stimulation.

**Legend Movie 9**
Ionomycin treatment of cells expressing APPL1 GFP and RGECO.

**Legend Movie 10**
EGF stimulation of cells expressing APPL1 GFP and RGECO.
